# Authentication of In Situ Measurements for Thoracic Aortic Aneurysms in Mice

**DOI:** 10.1101/2020.12.24.424013

**Authors:** Satoko Ohno-Urabe, Masayoshi Kukida, Michael K. Franklin, Yuriko Katsumata, Wen Su, Ming C. Gong, Hong S. Lu, Alan Daugherty, Hisashi Sawada

## Abstract

Aortic diameter is a standard parameter for defining disease severity of thoracic aortic aneurysms. In mouse studies, aortic diameters can be measured in situ directly, but this approach has a potential confounder of underestimation due to the absence of physiological arterial blood pressure. In the present study, we developed an in situ approach for authentic aortic measurements. Thoracic aortic aneurysms were induced by beta-aminopropionitrile (BAPN, 0.5% wt/vol) administration in 4-week-old male C57BL/6J mice. Ultrasonography was performed to examine aortic dimensions, and mice with thoracic aortic dilatations were terminated subsequently. After saline perfusion through the left ventricle, periaortic tissues were removed to expose thoracic aortas. Optimal cutting temperature (OCT) compound was injected via the left ventricle to maintain aortic patency. In situ aortic images were captured pre- and post-OCT injection. In mice with severe thoracic aortic aneurysms, smaller aortic diameters were observed prior to OCT injection compared to ultrasound measurements, while aortic diameters in situ after OCT were comparable to diameters measured using ultrasound. A telemetry system demonstrated that maximal luminal pressures during 150 ul of OCT injection were 90 mmHg. Immunostaining for CD31 revealed that endothelial cells were preserved in the intima after OCT injection. These results indicate that OCT injection does not cause aortic damages due to excess pressures. In conclusion, in situ imaging with OCT injection provides authentic aortic measurements without overt aortic damage in mice with thoracic aortic aneurysms.

---

Aortic dimension is the most commonly used criterion of the severity of thoracic aortic aneurysms and can be determined by several approaches in mice.^1^ In situ imaging has been used to measure aortic diameters.^2^ However, a potential caveat of aortic measurements using this mode is the absence of arterial blood pressure that can lead to under-estimation of aortic dimeters, particularly in mice with a flaccid aortic wall. The present study developed an in situ imaging approach to overcome this shortcoming, and demonstrated authentic dimensions of thoracic aortic aneurysms in mice.

β-aminopropionitrile (BAPN, 0.5% wt/vol in drinking water) was administered to induce thoracic aortic aneurysms in C57BL/6J mice (male, 4-week-old, n=30).^3^ Ultrasound imaging (Vevo 2100, MS550) was performed after 4, 8, and 12 weeks of BAPN administration, and aortic diameters were measured at the most dilated area at end-diastole. During BAPN administration, 16 mice died and ultrasonography detected a wide range of luminal dilatations of the thoracic aorta from mild to severe in 10 mice (**Figure A, B, Supplemental Figure 1**). In these 10 mice, 7 mice were randomly selected and in situ imaging was performed. The right atrial appendage was excised and saline (8 ml) was perfused through the left ventricle. Perivascular tissues of the thoracic aorta were removed gently and a black plastic sheet was inserted behind the aorta to enhance contrast of the aortic wall. Optimal cutting temperature (OCT) compound (150 µl) was then introduced into the left ventricle in 30 seconds using an insulin syringe to maintain aortic patency. Two dimensional images and cine loops were recorded from pre-injection to 50 seconds post-OCT injection using a dissection microscope with a high-resolution camera (#SMZ800, #DS-Ri1, Nikon). Imaging procedure was completed within 10-15 minutes in each mouse. To determine the aortic patency during imaging, aortic diameters were compared between ultrasound and in situ images. Several mice exhibited considerably smaller aortic diameters in images acquired prior to introduction of OCT compared to diameters measured using ultrasound (−0.8 to −0.4 mm), and OCT injection inflated the aortic dimensions (**Figure A, B, Supplemental Video 1**). Bland-Altman plots revealed that the bias between in situ and ultrasound diameters was closer to 0 in post-than that in pre-OCT images (**Figure C**). The range of limits of agreement was smaller in post-OCT (−0.4 to 0.3 mm) than in pre-OCT (−1.0 to 0.5 mm). Furthermore, a significant positive correlation between ultrasound and in situ measurements was only observed in post-, but not pre-, OCT images (**Figure D**). Therefore, in situ imaging using OCT injection provided more accurate aortic measurements in mice with thoracic aortic aneurysms.

**Figure A.**
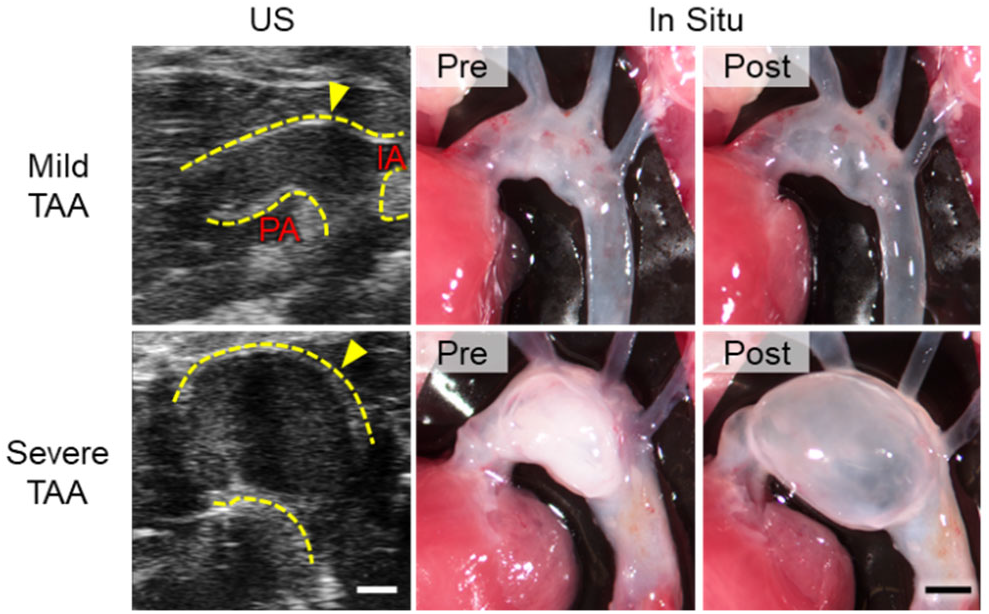
Representative ultrasound and in situ images of thoracic aortas in BAPN-administered mice. Post-OCT images were captured 50 seconds after completion of OCT injection. TAA indicates thoracic aortic aneurysm; Yellow dotted lines, the aortic wall; IA, innominate artery; PA, pulmonary artery. Scale bar, 1 mm. n=7.

**Figure B.**
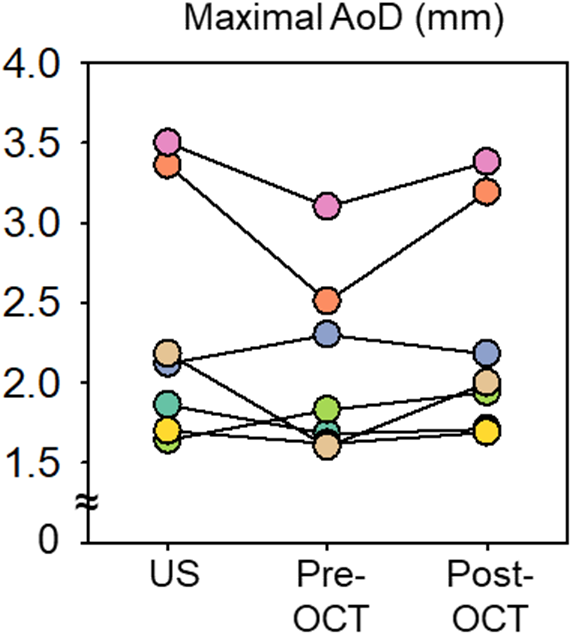
Aortic diameters measured at the most dilated area of ultrasound and in situ images with/without OCT injection. n=7.

**Figure C.**
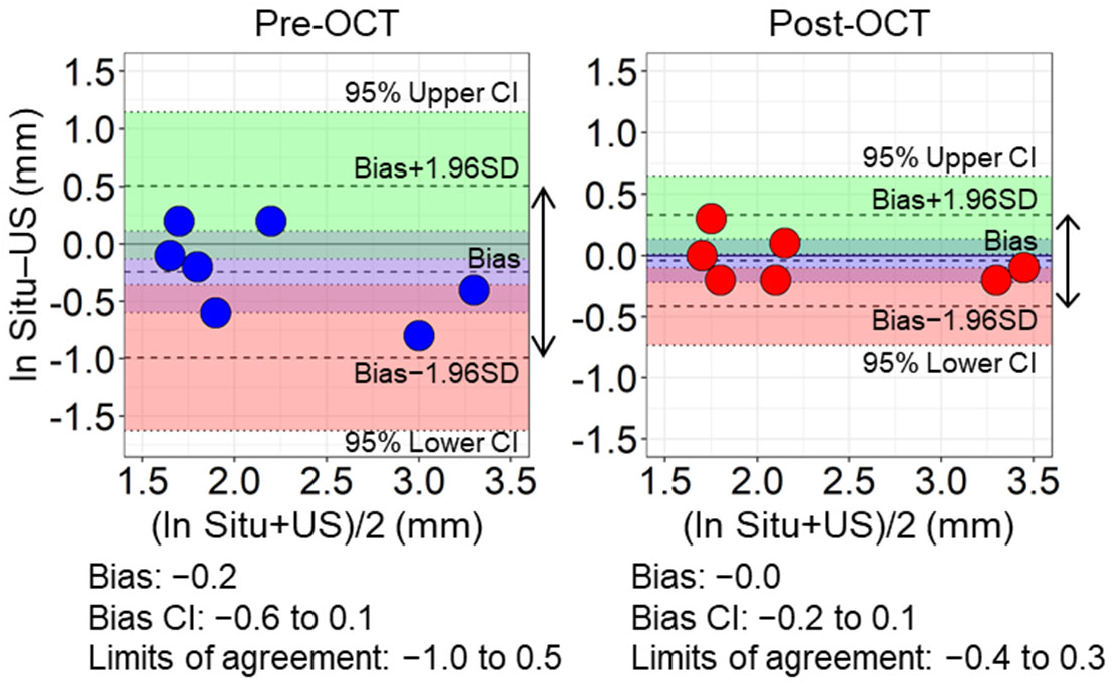
Bland-Altman plots reveal the low variability of aortic measurements after OCT injection. Double arrows indicate limits of agreement. Light green and red areas; 95% lower and upper limit agreement confidence intervals.

**Figure D.**
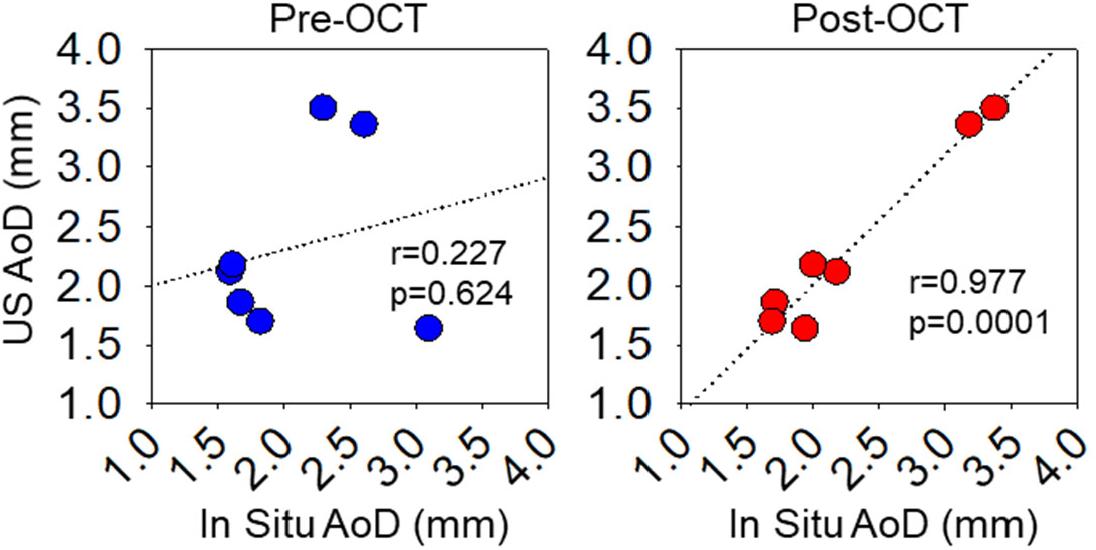
Correlations of aortic measurements between ultrasound and in situ pre- or post-OCT images.

OCT was injected using a small gauge needle (28G) that provided a slow flow of this viscous material. We measured luminal pressures during OCT injection using a telemetry system (#TA11PA-C10, Data Sciences International) in C57BL/6J male mice (n=5). Luminal pressures were 90±18 mmHg at the peak and decreased to less than 50 mmHg after at 10 seconds (**Figure E**). Then, aortic diameters were measured during OCT injection in mice with severe aortic dilatation (n=3). Aortic diameters were the largest immediately following OCT injection, with a subsequent modest reduction that was plateaued after at 40 seconds (**Figure F, Supplemental Video 2**). Importantly, maximal external diameters measured with in situ images were comparable to luminal diameters at mid systole in ultrasound images (ultrasound: 3.0±0.4; in situ: 3.0±0.4 mm, p=0.70). Endothelial cells were also examined by immunostaining for CD31 to determine whether OCT injection led to loss of endothelial cells. Aortic tissues were harvested from C57BL/6J male mice after OCT injection (n=5). CD31 positive cells were detected throughout the intima of both ascending and descending thoracic aortas (**Figure G**). These results demonstrated that OCT injection did not cause aortic over-expansion due to pressure overload and overt aortic tissue damage.

High frequency ultrasound is a powerful tool to evaluate aortic diameters in mice, and have been used by many studies.^4, 5^ However, the availability of ultrasound systems is restricted by its expense. Also, ultrasonography requires appropriate training for reliable imaging and authentic measurements. In addition, some mouse models of thoracic aortic aneurysms involve the descending aorta which is not readily imaged by ultrasonography. The present study demonstrated in situ image with OCT injection exhibited comparable aortic measurements to ultrasonography. Furthermore, in situ images can measure aortic dimensions directly in any regions of the thoracic aorta. Thus, in situ imaging with OCT injection can be considered as not only an optimal alternative approach but also a validation mode to ultrasound aortic measurements. The combination of ultrasound and in situ measurements would provide more robust evaluation of the severity of thoracic aortic aneurysms in mice.

**Figure E.**
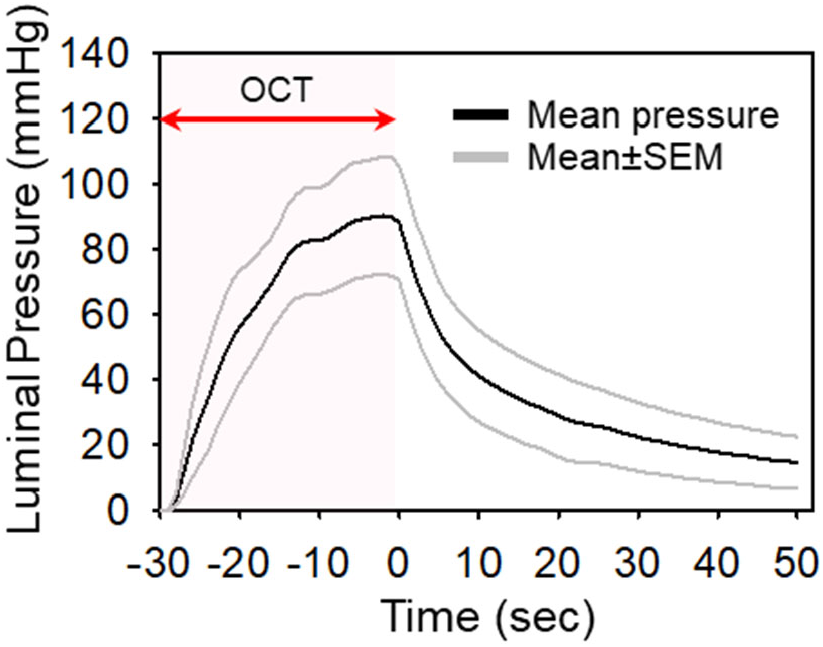
Luminal pressures during OCT injection in normal aortas (n=5).

**Figure F.**
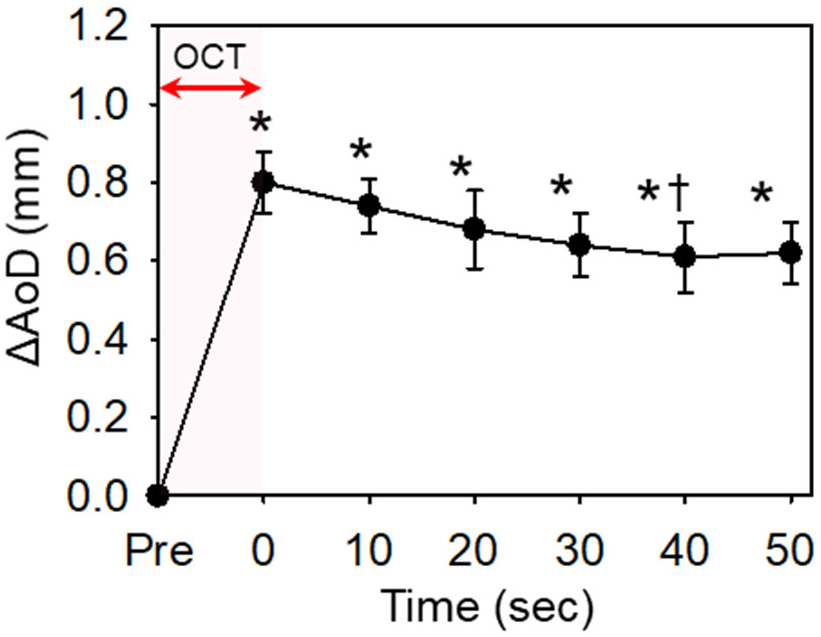
Sequential transition of aortic diameters after OCT injection in aneurysmal tissues (n=3). * p<0.05 vs Pre, † p<0.05 vs 0 sec by One-Way repeated measures ANOVA followed by t-test with Bonferroni correction.

**Figure G.**
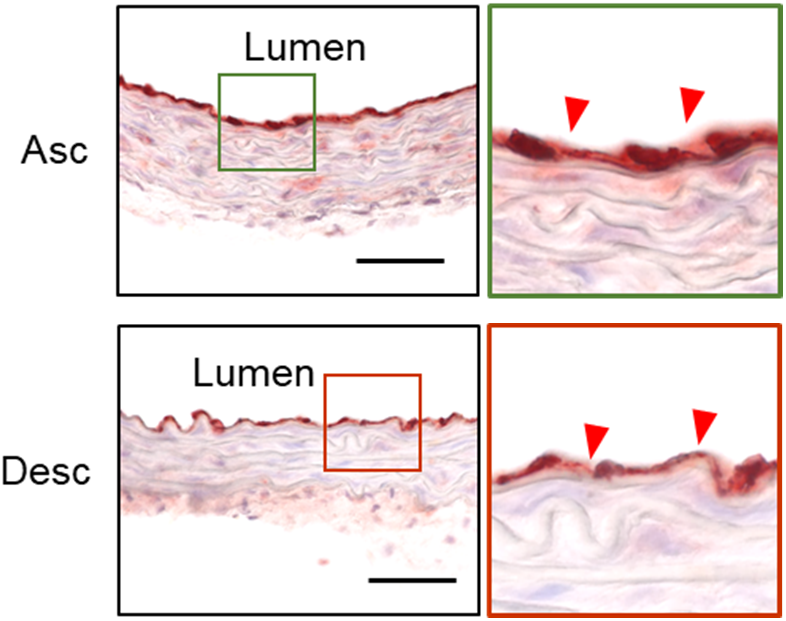
Immunostaining for CD31 of normal aortas with OCT injection. n=5. Red triangles indicate endothelial cells; L, lumen; Asc, ascending aorta; Desc, descending; scale bar, 50 µm.

Aortic diameters can be measured by ex vivo approach; however, aortic patency is often not maintained during ex vivo imaging of aneurysmal aortas, which may lead to underestimation of aortic dimensions. Therefore, formalin or latex perfusion are frequently utilized to recapitulate aortic morphology ex vivo.^2,6, 7^ However, these procedures need a perfusion system and are time-consuming. OCT is a nonhazardous reagent and can be readily applied. In addition, OCT did not cause discernable tissue damage. Therefore, OCT injection is a simple and effective procedure for in situ aortic measurements. Notable, in situ images measure aortic diameters in a lateral (right-left) axis. Imaging planes should be optimized individually, especially mice with significant aortic dilatation on the anterior or posterior wall.

In conclusion, OCT injection is an optimal approach for maintaining aortic patency in situ, which provides more reliable aortic measurements in mice.

## Supporting information

Raw data

## Materials and Methods

All raw data and analytical methods are available from the corresponding author upon appropriate request.

### Mice

C57BL/6J male mice at 3 weeks of age were purchased from The Jackson Laboratory (Stock# 000664, Bar Harbor, ME). Mouse housing conditions are described in **Table 1**. Mice were housed 5 per cage and allowed access to diet (Teklad Irradiated Global 18% Protein Rodent Diet, Cat# 2918, Envigo, Madison, WI) and water *ad libitum*. Bedding was changed weekly during the study, and cotton pads were provided as enrichment. The room was maintained at a 14:10 hour light:dark cycle, constant temperature of 18-23 °C, and 40-60% humidity. All protocols were approved by the University of Kentucky IACUC.

**Table 1.** Mouse Housing Conditions.

| Parameter | Conditions | Note |
| --- | --- | --- |
| Housing density | 5 per cage |  |
| Water | 0.5% BAPN mixed Water | ad libitum |
| Diet | Teklad Irradiated Global 18% Protein Rodent Diet, Cat# 2918; Envigo | ad libitum |
| Bedding | P.J. Murphy Coarse SaniChip<br>Cotton pads for enrichment |  |
| Light Cycle (light:dark) | 14:10 hours |  |
| Set Temperature range | 18-23 °C |  |
| Set Humidity range | 40-60% |  |

### Induction of thoracic aortic aneurysms

β-aminopropionitrile (0.5% wt/vol, Cat# A3134, Millipore-Sigma, St. Louis, MO) was administered through drinking water at 4 weeks of age.^3^ BAPN drinking water was replaced twice a week.

### In vivo and in situ evaluation of thoracic aortic aneurysms

Thoracic aortas were scanned at 4, 8, and 12 weeks of BAPN administration using a Vevo 2100 ultrasound system with a MS 550 transducer (FUJIFILM VisualSonics Inc., Canada). During ultrasound scanning, mice were placed on a heated platform (37°C) to avoid hypothermia and anesthetized by isoflurane (1-2% vol/vol) to adjust the heart rate between 400 to 550 beats/minutes. Longitudinal images of ascending and descending aortas were captured in the parasternal or paraspinal approach respectively, as described previously.^4,5^ Aortic diameters were measured at end diastole in 3 different cardiac cycles. Ultrasound measurements were performed using Vevo LAB 3.1.1 software (FUJIFILM VisualSonics Inc.)

Mice in which ultrasound detected aortic dilatation were euthanized at the same day or one day after ultrasonography by ketamine:xylazine cocktail (90, 10 mg/kg, respectively). The right atrial appendage was excised and saline (8 ml) was perfused via the left ventricle. Periaortic tissues were removed gently and a black plastic sheet was then inserted under the aorta. Thoracic aortas were inflated by injection of optimum cutting temperature compound (150 µl, OCT, Cat# 23-730-571, Thermo Fisher Scientific, Waltham, MA) from the left ventricle using an insulin syringe with 28-gauge needle. In situ aortic images were captured before, and 50 seconds after, OCT injection with a Nikon SMZ800 stereoscope (Nikon, Tokyo, Japan). Cine loops were also recorded during OCT injection. All aortic in situ images were captured with a small ruler placed next to the aorta. Aortic images were analyzed using NIS-Elements AR software (Ver 4.51, Nikon). The measurement software was calibrated using the ruler on individual images. To measure aortic diameters, a measurement line was drawn perpendicularly to the aortic axis at the most dilated area. Aortic measurements were verified by an independent investigator who was blinded to the initial analysis.

### Measurements of luminal pressures during OCT injection

Luminal pressures were measured using a telemetry system (#TA11PA-C10, Data Science International, St. Paul, MN) in 10-12-week-old C57BL/6 mice (n=5). Mice were euthanized by the ketamine:xylazine cocktail followed by saline perfusion. Periaortic tissues were removed to expose aorta in situ. Subsequently, a telemetry transducer was inserted into the left common carotid artery with the tip located at the branching point of the aortic arch. OCT was injected from the apex of left ventricle (150 µl in 30 seconds). Luminal pressures were recorded from the start of OCT injection to 50 seconds after the completion of OCT injection.

### Immunostaining

Ascending and descending aortas were harvested and fixed with paraformaldehyde (4% wt/vol) overnight followed by 24 hours incubation with sucrose (30% wt/vol). Aortic samples were then embedded in OCT and cut into 10 µm sections. Slides were baked at 60 °C for 20 minutes and immersed in H_2_O_2_ (1% vol/vol) in methanol to quench endogenous peroxidase. Sections were incubated with normal goat serum (2.5% vol/vol) for 20 minutes. Rat anti mouse CD31 (1 µg/ml, Cat# ab7388, abcam, Cambridge, MA) was used as the primary antibody. Detection of the primary antibody was visualized using Rat ImmPRESS (Cat# MP-7444, Vector Laboratories, Burlingame, CA) and ImmPACT AEC kits (Cat# SK-4205, Vector Laboratories). Tissue sections were counterstained subsequently with hematoxylin. Histological images were captured using an Eclipse E600 microscope with a DSRi1 digital camera (Nikon).

### Statistical analyses

SigmaPlot 14.0 (SYSTAT Software Inc., Palo Alto, CA) was used for statistical analyses. All data were expressed as mean±standard error of the mean. Normality and homogeneous variance were confirmed in all data by Shapiro-Wilk and Brown-Forsythe tests, respectively. Correlations between in vivo and in situ aortic diameters was evaluated by Pearson correlation coefficient. The transition of aortic diameters during OCT injection was examined by one-way repeated measures ANOVA after square root transformation. Bland-Altman plots were generated by R (version 3.6.3) using the blandr package (version 0.5.1).^8^ p<0.05 was considered statistically significant.

