## Supplemental Figure for "Authentication of In Situ Measurements for Thoracic Aortic Aneurysms in Mice"

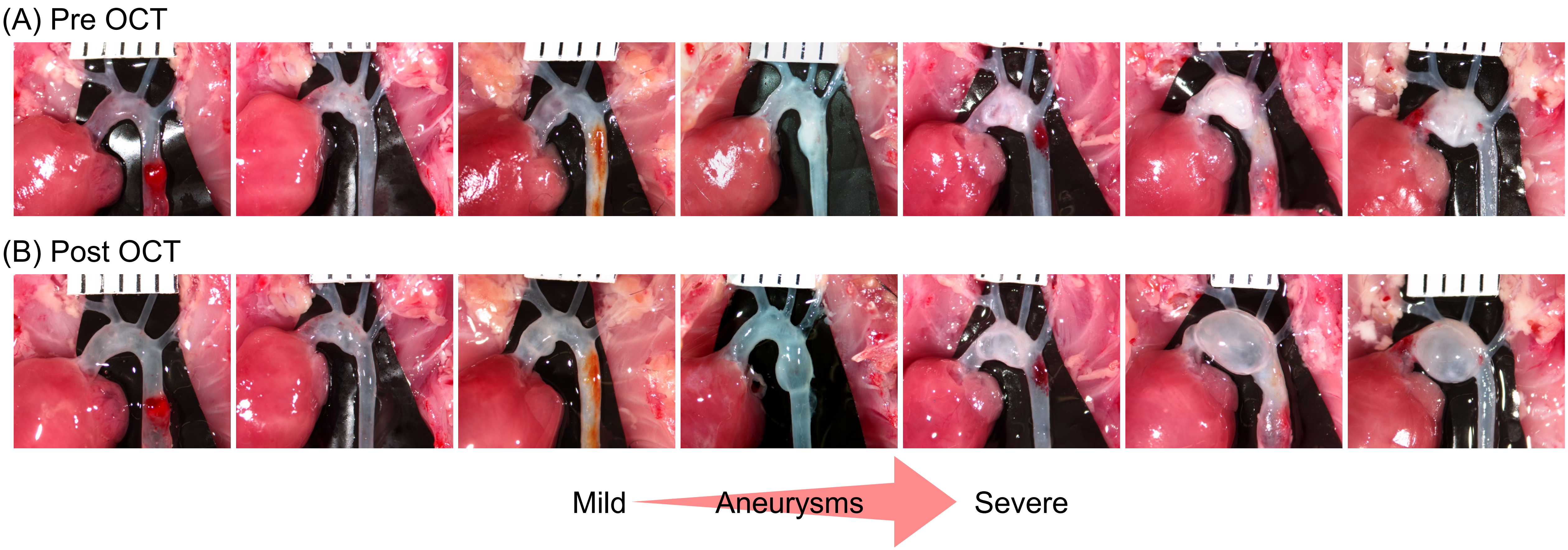


**Supplemental Figure.** In situ images of **(A)** pre- and **(B)** post-OCT injection for thoracic aortic aneurysms in BAPN-administered mice. Images are placed in order of the severity of thoracic aortic aneurysms from mild to severe.
